# Serine and glycine are essential for human muscle progenitor cell population expansion

**DOI:** 10.1101/833798

**Authors:** Brandon J. Gheller, Jamie E. Blum, Erica L. Bender, Mary E. Gheller, Esther W. Lim, Michal K. Handzlik, Patrick J. Stover, Martha S. Field, Benjamin D. Cosgrove, Christian M. Metallo, Anna E. Thalacker-Mercer

## Abstract

Skeletal muscle regeneration is reliant on a population of muscle specific adult stem cells (muscle progenitor cells; MPCs). During regeneration, the MPC population undergoes a transient and rapid period of population expansion, which is necessary to repair damaged myofibers and restore muscle homeostasis. Much research has focused on the age-related accumulation of negative regulators of regeneration, while the age-related decline of nutrient and metabolic determinants of the regenerative process needs examination. We hypothesized that older individuals, a population that is at risk for protein malnutrition, have diminished availability of amino acids that are necessary for MPC function. Here, we identified that levels of the non-essential amino acid serine are reduced in the skeletal muscle of healthy, older individuals. Furthermore, using stable-isotope tracing studies, we demonstrate that primary, human MPCs (*h*MPCs) exhibit a limited capacity for *de novo* biosynthesis of serine and the closely related amino acid glycine. We identified that serine and glycine are essential for *h*MPC proliferation and, therefore, population expansion. Serine and glycine were necessary to support synthesis of the intracellular antioxidant glutathione, and restriction of serine and glycine was sensed in an EIF2α-dependent manner resulting in cell cycle arrest in G0/G1. In conclusion, we elucidate that, despite an absolute requirement of serine/glycine for *h*MPC proliferation, availability of serine in the skeletal muscle microenvironment is limited to the *h*MPCs of healthy older adults and is a likely underlying mechanism for impaired skeletal muscle regeneration with advancing age.

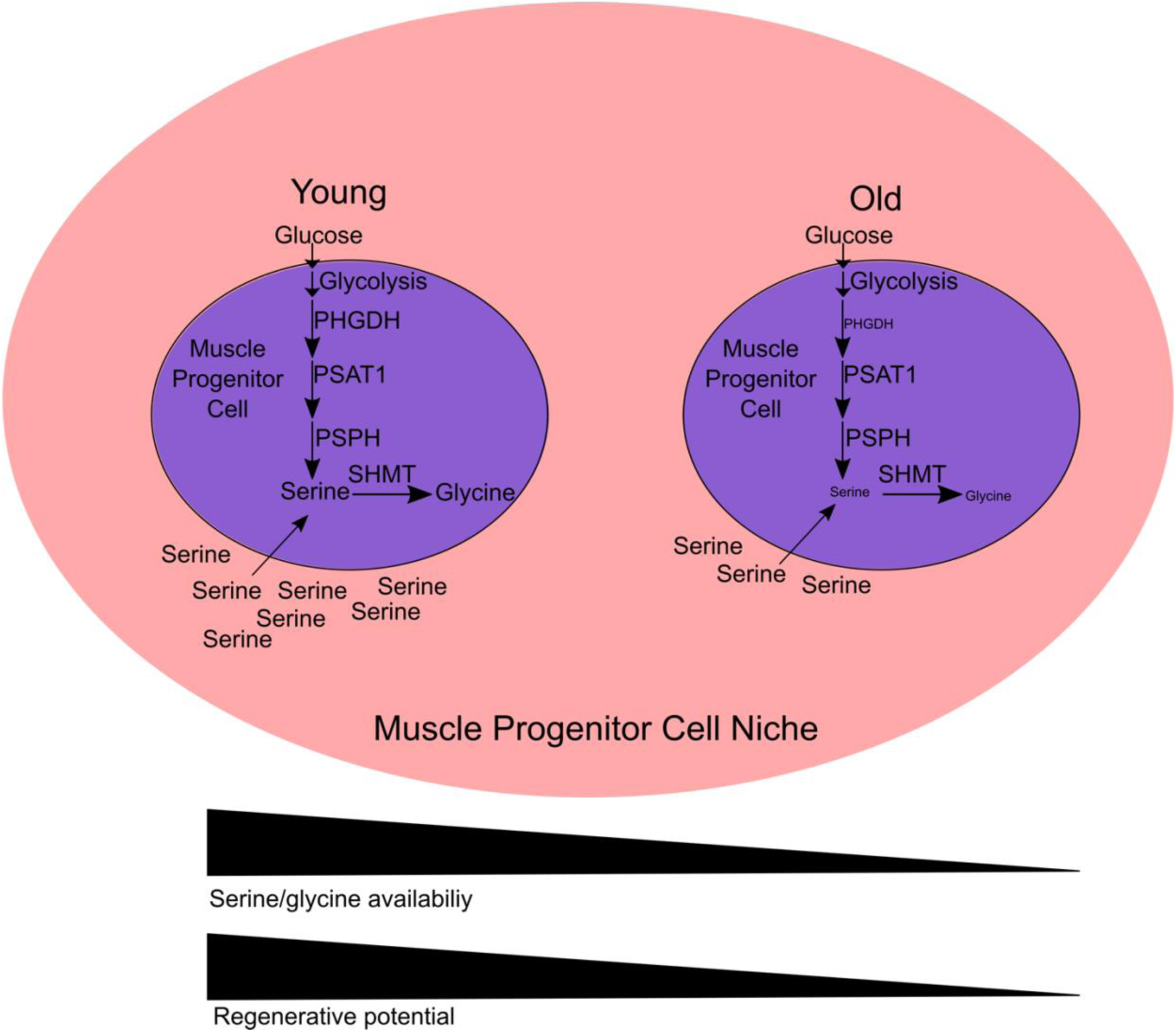
Graphical Abstract.

## Introduction

Skeletal muscle regeneration is reliant on a population of muscle specific adult stem/ progenitor cells (MPCs) identified by the canonical transcription factor PAX7 (Seale et al., 2000). MPCs reside in a quiescent state, are activated after injury, and undergo a transient and rapid population expansion to provide an adequate number of cells to donate nuclei to damaged myofibers or to create nascent myofibers thereby restoring homeostasis. The muscle regeneration process markedly declines with age due to a myriad of MPC-intrinsic and -extrinsic factors that remain to be fully elucidated (Blau et al., 2015). Parabiosis experiments provide evidence that there are alterations in circulating factors and, ultimately the MPC microenvironment that occurs in aged animals which negatively affects muscle regeneration (Conboy et al., 2005). For example, when the circulation of a young and an aged mouse are joined, muscle regeneration after injury is improved in old mice compared to when the circulation of two old mice are joined (Conboy et al., 2005). Thus, restoration of the MPC microenvironment, to match that of young animals, is a potential avenue for improving skeletal muscle regeneration (Conboy et al., 2005). Follow-up to these landmark studies have largely focused on mechanisms downstream of ligand-based signaling pathways such as the Notch (Conboy et al., 2003) and Wnt (Brack et al., 2007) pathways as well as transforming growth factor beta pathways (Egerman et al., 2015; Sinha et al., 2014). Relatively little attention has focused on nutrient needs and availability to support MPC expansion. Intracellular amino acid availability has been shown to be affected by age in both a cell-intrinsic (e.g., alterations in metabolism) and -extrinsic (e.g., availability in circulation) manner (Dunn et al., 2014; Menni et al., 2013). For example, while gastrointestinal absorption of amino acids after feeding does not appear to be impaired with age (Katsanos et al., 2006; Mitchell et al., 2015), the profile of circulating amino acids is altered by age with lower levels of serine, alanine, proline, tyrosine, and methionine observed in older individuals (Houtkooper et al., 2011). The importance of amino acid availability to regenerative processes is highlighted by the observation that amino acids are the primary contributors to cell mass during proliferation (Hosios et al., 2016).

It has been previously documented that rapid changes in MPC metabolism occur after activation (Rodgers et al., 2014), likely to support metabolic demands associated with cell division (Hosios et al., 2016). These dynamic, metabolic changes in MPCs are likely accompanied by changes in exogenous nutrient requirements, particularly those traditionally considered non-essential, as demonstrated in other proliferative cell types (Ma et al., 2017). Elucidating the metabolic requirements of MPC proliferation is essential for optimizing muscle regeneration after injury, particularly in populations with impaired regeneration.

We aimed to determine if an age-related decline in amino acid availability exists in the human MPC (*h*MPC) microenvironment and to determine the impact of this amino acid decline on MPC activity if it exists. In this study we identify that serine is the only amino acid that declines in *h*MPC microenvironment with advancing age, and *h*MPCs possess a limited capacity for *de novo* biosynthesis of serine and the closely related amino acid, glycine. Without exogenous serine/glycine, *h*MPCs halt proliferation and arrest in a G0/G1 state. We attribute the requirement for exogenous serine/glycine in part to the need for glutathione synthesis and determined that the G0/G1 cell cycle arrest that occurs in response to serine/glycine restriction occurs in an EIF2α-dependent manner.

## Results

### Serine availability and metabolism is dysregulated in aged skeletal muscle

Analysis of human skeletal muscle biopsy tissue identified serine as the only measured amino acid that was reduced in the skeletal muscle of older adults compared to younger adults (**Figure 1A**). Further, chronological age and skeletal muscle serine levels were negatively correlated (**Figure 1B**). This finding is supported by a previous cross-sectional analysis of plasma from healthy, younger and older individuals which demonstrated that serine levels are reduced in older humans (Kouchiwa et al., 2012). Others have demonstrated that in frail older adults, skeletal muscle concentrations of serine and also glycine are reduced (Fazelzadeh et al., 2016). Coupled with lower circulating levels, decreased skeletal muscle serine levels suggest that after injury and myofiber disruption the *h*MPC microenvironment of aged individuals has less serine and potentially less glycine available.

**Figure 1.**
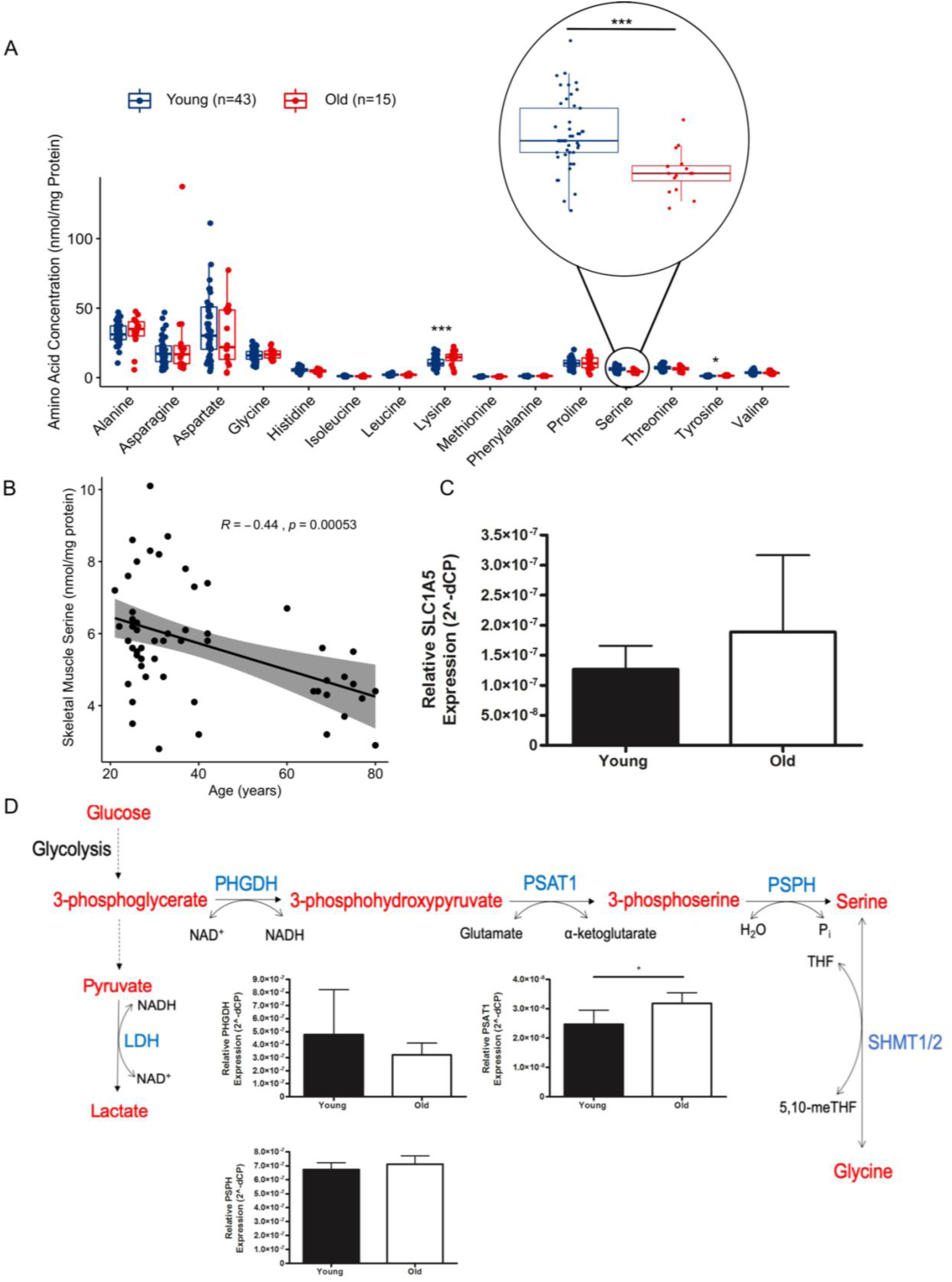
Serine availability and metabolism are dysregulated in aged skeletal muscle and *h*MPCs. A) Analysis of amino acid levels in skeletal muscle biopsy tissue from younger (20-45y, n=43) and older (60-80y, n=15) donors identify serine as the only amino acid that decreases with age (P<0.001). The normalcy of the distribution of each amino acid was assessed by the Shapiro-Wilk test. If data were determined to be normally distributed, they were compared via an unpaired t-test otherwise they were compared Mann-Whitney U-test. B) Skeletal muscle serine levels are negatively correlated with age (n=58, r=-.44, P<0.0001) based on a Pearson correlation. C) Gene expression of the serine transporter SLC1A5 (P>0.05) between skeletal muscle tissue from younger (n=11) and older donors (n=10). Data expressed as mean ± SD. D) A schematic of the serine/glycine biosynthesis pathway. Intermediary metabolites in red. Key enzymes in blue. PHGDH, phosphoglycerate dehydrogenase; PSAT1, phosphoserine amino transferase 1; PSPH, phosphoserine phosphatase; SHMT 1/2, serine hydroxy methyltransferase1/2; LDH, lactate dehydrogenase. Gene expression of PHGDH (P>0.05), PSAT1 (P<0.05), and PSPH (P>0.05) between skeletal muscle tissue from younger (n=11) and older donors (n=10). Data expressed as mean ± SD. *P<0.05.

To determine what limits serine levels in older skeletal muscle, we measured the gene expression of the serine transporter *SLC1A5* in skeletal muscle tissue homogenates from younger and older adults and found no age-related differences in expression (**Figure 1C**). We next measured gene expression of the serine synthesis enzymes, *PHGDH*, *PSAT1*, and *PSPH* in skeletal muscle tissue homogenates. *PSAT1* was the only gene that was differentially expressed in skeletal muscle with age (1.4-fold greater in older vs. younger, **Figure 1D**). Thus, it is unclear whether impaired uptake or biosynthesis limits serine levels in old muscle tissue.

### Serine and glycine are required for hMPC population expansion

Because skeletal muscle serine levels and the regenerative potential of *h*MPCs decline with advancing age, and because serine has previously been shown to be the third most consumed metabolite by proliferating mammalian cells (Hosios et al., 2016) we evaluated whether serine and/or glycine impact *h*MPC population expansion. Serine and glycine were considered alone or in combination due to their interconversion through one enzymatic step via the serine hydroxymethyltransferase (SHMT, **Figure 1D**). When *h*MPCs were cultured without serine and glycine, they did not undergo population expansion (**Figure 2A**). The addition of serine (**Figure 2B**) and glycine (**Figure 2C**) individually or in combination (**Figure 2A**) increased *h*MPC population expansion in a dose-dependent manner. At high doses, glycine alone was more effective at increasing *h*MPC population expansion than serine (**Figure 2D**). This is dissimilar to what is observed in other cell types in which serine and not glycine is required for population expansion (Labuschagne et al., 2014; Ma et al., 2017). We verified that serine/glycine restriction in the media reduced intracellular serine and glycine levels (**Figure 2E**).

**Figure 2.**
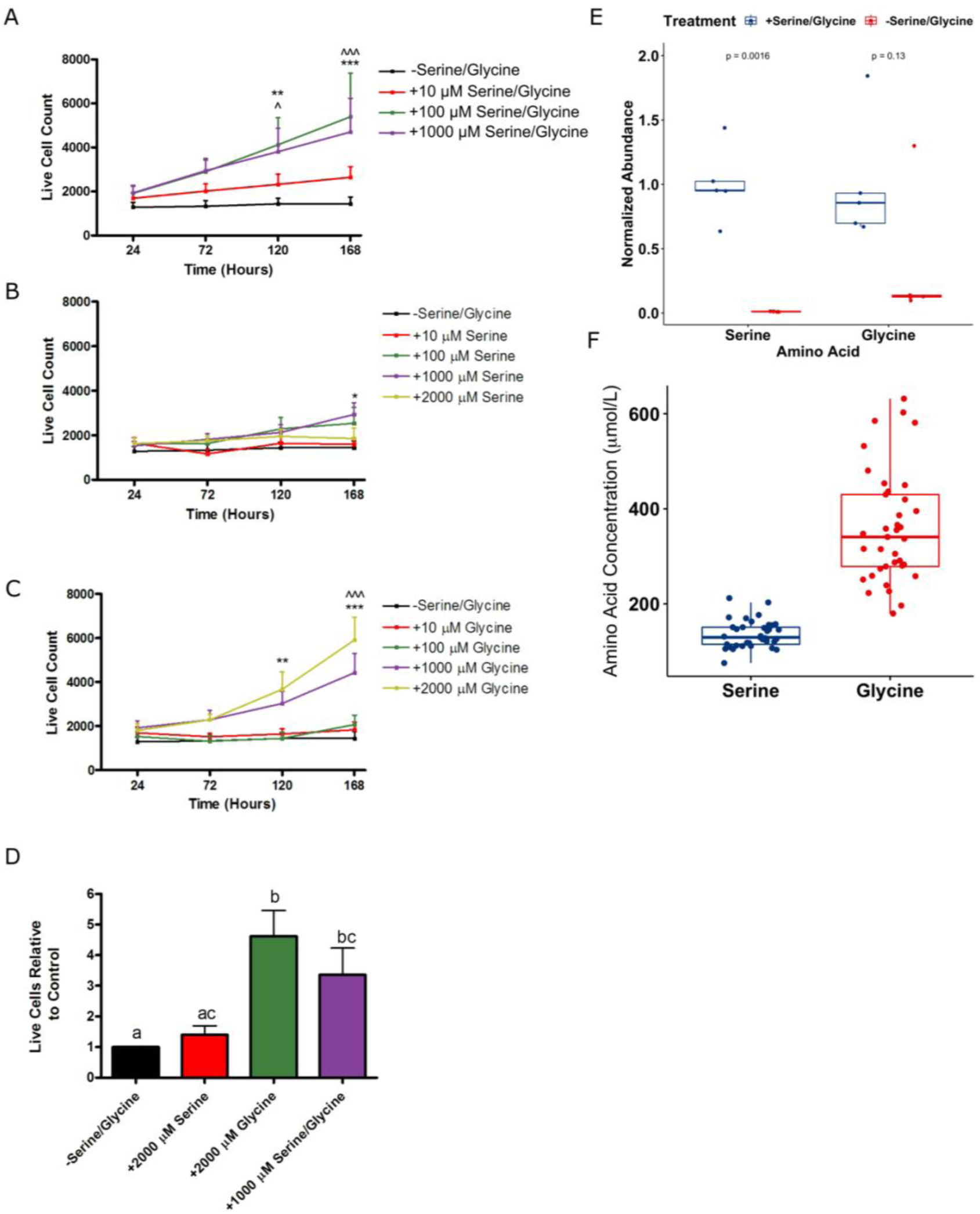
Serine and glycine are essential for *h*MPC population expansion. Live cell count was determined by co-staining cells with Hoescsht 33342 (to identify all cells) and propidium iodide (to identify dead cells) after *h*MPCs were cultured in media lacking serine/glycine or increasing concentrations of A) serine/glycine (**indicates significant difference between 100 µM vs. -serine/glycine, P<0.01, ***indicates significant difference between 100 µM vs. -serine/glycine, P<0.001, ^indicates significant difference between 1000 µM vs. -serine/glycine, P<0.05, ^^^indicates significant difference between 1000 µM vs. -serine/glycine, P<0.001), B) serine alone (*indicates significant difference between 1000µM vs. -serine/glycine, P<0.05), or C) glycine alone (**indicates significant difference between 2000 µM vs. -serine/glycine, P<0.01, ***indicates significant difference between 2000 µM vs. -serine/glycine, P<0.001, ^^^indicates significant difference between 1000 µM vs. -serine/glycine, P<0.001) Data are expressed as mean ± SD. D) The relative number of live cells after seven days of culture with the indicated concentrations of serine, glycine, or serine and glycine. Different letters are significantly different from each other. E) Levels of serine and glycine after 7 days of culture in serine/glycine restricted media as determined by GC-MS. F) Levels of serine and glycine in the plasma of fasted, young humans (n=37) based on GC-MS measurements.

To determine the physiological relevance of the serine and glycine concentrations used *in vitro*, we measured plasma serine and glycine concentrations from humans under fasting, resting conditions. The levels of plasma serine were between 76 and 212 µmol/L and plasma glycine were between 178 and 632 µmol/L (**Figure 2F**). However, the levels of these amino acids in circulation are frequently elevated above basal values in the non-fasted state (Gannon et al., 2002; Garofalo et al., 2011) Further, it is likely, that the localized, skeletal muscle availability is also a contributing factor. The physiological relevance of the concentrations used for cell culture experiments are also supported by an analysis by Bergström *et al*. demonstrating that in younger adults, skeletal muscle concentrations of serine and glycine are ∼980 μmol/L and ∼1330 μmol/L, respectively (Bergström et al., 2017).

### Serine/glycine restriction causes cell cycle arrest

We next sought to understand what drives differences in *h*MPC population expansion when *h*MPCs are cultured with and without serine/glycine. We observed that differences in population expansion, during serine/glycine restriction, could not be explained by cell death (**Figure 3A**). Using DNA staining and analysis via flow cytometry, we observed a greater number of cells in the early phases of the cell cycle with serine/glycine restriction (**Figure 3B**). This finding was supported by reduced incorporation of BrdU into DNA after a 24-hour pulse (**Figure 3C**) and an accumulation of CYCLIN D1 protein in *h*MPCs cultured in serine/glycine restricted media (**Figure 3D**). Because *h*MPC differentiation is modeled *in vitro* by serum restriction (Pavlidou et al., 2017), we measured the *h*MPC specific, mid-to late-differentiation marker MYOGENIN, which was undetectable (**Figure 3E**). Additionally, levels of PAX7 and MYOD were significantly elevated in serine/glycine restricted *h*MPCs (**Figure 3E**). Elevated MYOD levels further supports cell cycle arrest in G1; MYOD protein levels are typically increased in *h*MPCs during G1 of the cell cycle (Kitzmann et al., 1998).

**Figure 3.**
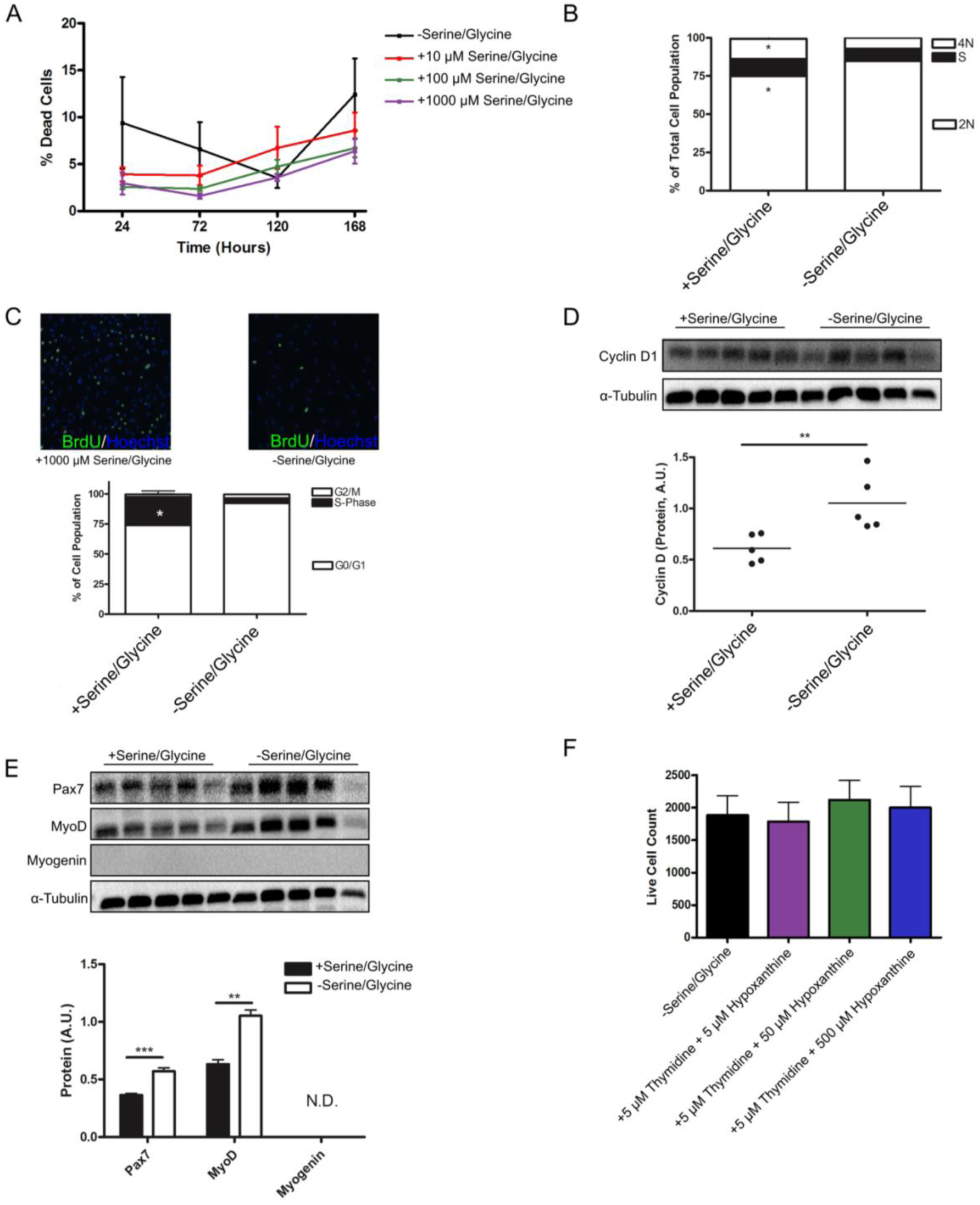
Serine/glycine restriction causes cell cycle arrest in G0/G1 in *h*MPCs. A) Percentage of dead cells was quantified by dividing all *h*MPCs which stained positive for propidium iodide by all *h*MPCs which stained positive for Hoechst 33342 after 5 days of culture in media with varying concentrations of serine/glycine. B) Propidium iodide staining and analysis via flow cytometry was used to determine DNA content of *h*MPCs after 5 days of serine/glycine restriction. *P<0.05. C) BrdU incorporation in after a 24-hour pulse in *h*MPCs undergoing serine/glycine restriction for 5 days. Data are expressed as mean ± SD. D) Immunoblot for CYCLIN D1 protein normalized to α-TUBULIN for quantification in *h*MPCs that had been serine/glycine restricted for 5 days. **P<0.01. E) Immunoblot for PAX7, MYOD, and MYOGENIN protein normalized to α-TUBULIN for quantification in *h*MPCs that had been serine/glycine restricted for 5 days. **P<0.01, ***P<0.001. N.D., not detectable. Data are expressed as mean ± SD. F) Live cell count for *h*MPCs treated with varying concentrations of thymidine and hypoxanthine. Data are expressed as mean ± SD. All experiments were repeated with *h*MPCs derived from the same 5 donors.

Previously, several proliferative cell types have demonstrated an exogenous requirement for serine/glycine (Labuschagne et al., 2014; Ma et al., 2017). The serine/glycine requirement in these cell types was attributed to DNA synthesis as the proliferative arrest can be rescued by glycine and formate, a one carbon unit integrated into the purine ring structure and necessary for the synthesis of thymidine, or by exogenous nucleotides. In contrast, in *h*MPCs the serine/glycine requirement appears to be derived from glycine (**Figure 2CD**) and is not DNA synthesis based as supplementation with nucleotide precursors did not rescue *h*MPC population expansion (**Figure 3F**).

### hMPCs exhibit limited capacity for serine/glycine biosynthesis

We hypothesized that *h*MPCs lack the ability for *de novo* serine/glycine biosynthesis, and that this underlies the exogenous requirements when *h*MPCs are cultured *in vitro*. In contrast to this hypothesis, RNA-seq analyses revealed that *h*MPCs drastically upregulate serine/glycine biosynthesis genes in response to serine/glycine restriction (**Figure 4A, Table S1**). We further verified that the enzymes directly implicated in serine/glycine biosynthesis (PHGDH, PSPH, PSAT1, SHMT1, SHMT2) are expressed at the protein level and the serine synthesis proteins, specifically, are upregulated in response to serine/glycine restriction (**Figure 4B**). The serine synthesis pathway branches from glycolysis at 3-phosphoglycerate, using glucose as the initial substrate (**Figure 1D**); however, glucose uptake did not increase after serine/glycine restriction (**Figure 4C**). Further, increasing glucose concentrations in the media did not affect *h*MPC proliferation in the absence of serine/glycine (**Figure 4D**), which suggests a limited capacity for serine/glycine biosynthesis by *h*MPCs. Using stable isotope tracing with _13_C_6_ glucose, we determined that *de novo* biosynthesis accounts for no detectable intracellular serine and glycine when serine/glycine are available in the media (**Figure 4E**). During serine/glycine restriction, *de novo* biosynthesis contributes ∼25% of serine and ∼5% of glycine to the intracellular pools (**Figure 4E**) demonstrating that *h*MPCs have the capacity for serine/glycine biosynthesis under restricted conditions. However, the increase in *de novo* biosynthesis is ineffective to maintain the requirements for population expansion as highlighted by the low levels of relative serine and glycine even when *de novo* biosynthesis is active (**Figure 2E**). Even if *h*MPCs were able to synthesize sufficient serine and glycine to support proliferation, we identified that primary *h*MPCs from older adults have reduced expression of the serine biosynthesis enzyme PHGDH, (**Figure 4F**, n=10) suggesting that *h*MPC-intrinsic *de novo* serine synthesis is likely impaired.

**Figure 4.**
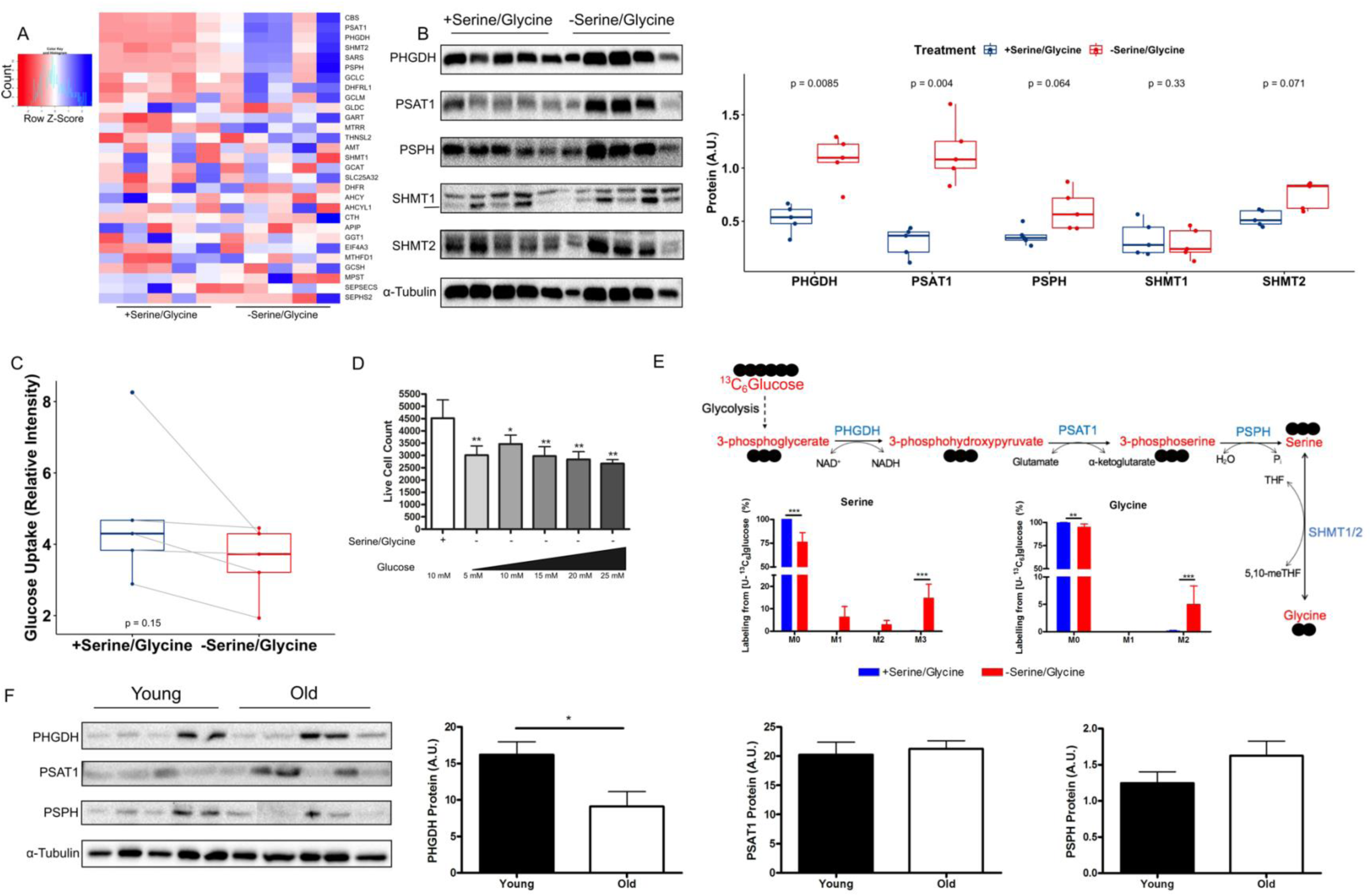
*h*MPCs exhibit limited capacity for serine/glycine biosynthesis. A) Heatmap of genes involved in serine, glycine, and one-carbon metabolism based on RNA-sequencing data from *h*MPCs after 5 days of culture in serine/glycine replete (1000 µM) or restricted conditions. B) Immunoblots for PHGDH, PSAT1, PSPH, SHMT1, and SHMT2 protein normalized to α-TUBULIN for quantification in *h*MPCs that had been serine/glycine restricted for 5 days. C) Glucose uptake by *h*MPCs after 5 days of culture in serine/glycine replete or restricted conditions. D) Live cell count of *h*MPCs cultured in serine/glycine replete media or serine/glycine restricted media and varying doses of glucose. *P<0.05, **P<0.01 relative to serine/glycine containing control. Data are expressed as mean ± SD. E) Percent mass isotopomer distribution of [U-_13_C]-glucose-derived serine and glycine in *h*MPCs cultured in serine/glycine replete (1000 µM) or restricted media for 5 days followed by 48 hours in similar media but containing [U-_13_C]-glucose. Data are expressed as mean ± SD. F) Protein levels of the serine/glycine biosynthesis enzymes PHGDH, PSAT1, and PSPH in *h*MPCs obtained from younger (n=5) and older (n=5) individuals as determined by immunoblotting. *P<0.05. Data expressed as mean ± SD. All experiments were repeated with *h*MPCs derived from the same 5 donors. P-values are indicated on the appropriate graphs.

### Serine/glycine restriction promotes oxidative stress in hMPCs and depletes intracellular glutathione

Several genes related to glutathione biosynthesis were upregulated with serine/glycine depletion in the transcriptomic dataset (**Figures 4A, Figure 5A, Table S1**). Because glycine is one of the three amino acids that comprise the tripeptide glutathione, we hypothesized that serine/glycine restriction reduced glutathione levels in *h*MPCs and impaired population expansion. Serine/glycine restriction decreased total glutathione levels by approximately 9-fold in *h*MPCs (**Figure 5B**) and decreased the ratio of GSH:GSSG (**Figure 5C**). As would be expected with lower glutathione levels, the primary intracellular antioxidant, we observed an increase in the levels of reactive oxygen species (ROS, **Figures 5D**). Supplementation with a cell permeable version of glutathione, glutathione ethyl ester (GSHee), modestly increased intracellular glutathione levels (**Figure 5E**) and decreased ROS levels (**Figure 5F**). Despite only a modest effect on intracellular glutathione levels, GSHee did provide a minor rescue to *h*MPC population expansion (**Figure 5G**). Of note, high levels of cell permeable glutathione were toxic (**Figure S1**) and therefore, we were not able to investigate how increasing intracellular levels of glutathione to that observed in serine/glycine replete cells would impact *h*MPC population expansion. We propose that ROS, associated with *h*MPC proliferation, cannot be counterbalanced during serine/glycine restriction due to reduced glutathione synthesis caused by limited serine/glycine synthesis leading to cell cycle arrest.

**Figure 5.**
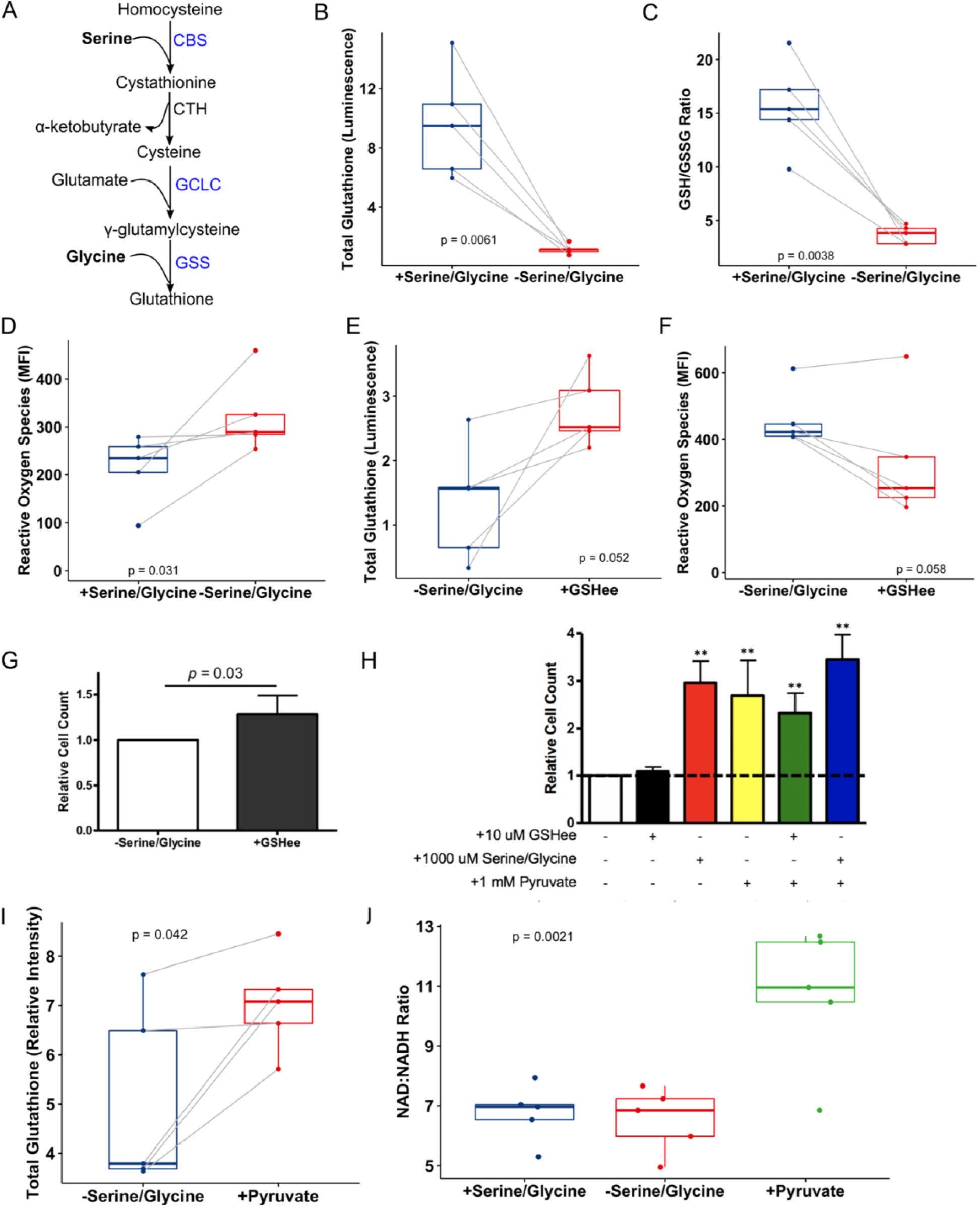
Serine/glycine restriction impairs glutathione synthesis but can be rescued by pyruvate supplementation. A) The glutathione synthesis pathway with genes upregulated after serine/glycine restriction according to RNA-sequencing in blue. CBS, cystathionine β-synthase; CTH, cystathionine gamma-lyase; GCLC, glutamate-cysteine ligase catalytic subunit; GSS, glutathione synthetase. B) Total intracellular glutathione levels in *h*MPCs after 5 days in serine/glycine restricted or replete media. C) Ratio of reduced to oxidized intracellular glutathione levels in *h*MPCs after 5 days in serine/glycine restricted or replete media. GSH, reduced glutathione; GSSG, oxidized glutathione. D) Reactive oxygen species in *h*MPCs after 5 days in serine/glycine restricted or replete media. E) Total intracellular glutathione levels in *h*MPCs after 5 days of serine/glycine restriction with or without 10 µM cell permeable glutathione ethyl ester (GSHee). F) Reactive oxygen species in *h*MPCs after 5 days of serine/glycine restriction with or without 10 µM GSHee. G) Live cell count after 5 days of serine/glycine restriction with or without GSHee (10 µM). H) Live cell count after 5 days of the indicated combination and concentration of serine/glycine, GSHee, and pyruvate. **P<0.01 I) Total glutathione levels in *h*MPCs cultured in a serine/glycine restricted media with or without 1 mM pyruvate for 5 days. J) Ratio of oxidized NAD (NAD_+_):reduced NAD (NADH) in *h*MPCs cultured in restricted serine/glycine media or media containing 1000 µM serine/glycine or 1 mM pyruvate. All experiments were repeated with *h*MPCs derived from the same 5 donors. P-values are indicated on the appropriate graphs.

### Pyruvate rescues hMPC proliferation during serine/glycine restriction

Under the assumption that inadequate glycine availability prevents adequate glutathione synthesis and ROS scavenging, we provided pyruvate in the absence of serine/glycine as an alternative antioxidant; we observed a complete rescue of the proliferative defect caused by serine/glycine restriction (**Figure 5H**). Pyruvate also increased total glutathione levels (**Figure 5I**). An additional mechanism by which pyruvate supplementation may have rescued *h*MPC proliferation, during serine/glycine restriction, is by increasing the NAD_+_/NADH ratio through the reduction of pyruvate to lactate, which provides the necessary NAD_+_ cofactor for serine biosynthesis (i.e., for the conversion of 3-phosphoglycerate to 3-phosphohydroxypyruvate, **Figure 1C**). Supporting this mechanism, pyruvate supplementation, in the absence of exogenous serine/glycine, increased the NAD_+_/NADH ratio which has recently been identified as the limiting cofactor for serine biosynthesis (Diehl et al., 2019) (**Figure 5J**).

### Serine/glycine restriction induces the integrated stress response in hMPCs

We hypothesized that ATF4 may integrate the sensing of serine/glycine restriction and underlie the transcriptional upregulation of the serine/glycine biosynthetic genes (**Figure 4A**). We noted several ATF4 target genes that were affected by serine/glycine restriction in *h*MPCs (**Figure 6A**). We further verified that protein levels of ATF4 and its upstream regulator, p-eIF2α, are increased after serine/glycine restriction (**Figure 6B**). Phosphorylation of eIF2α is known to coordinate a global decline in protein synthesis. In support of this, in the absence of serine/glycine, we observed an accumulation of most of the free, non-serine/glycine amino acids (**Figure 6C**) and a trend towards reduced protein synthesis (assessed by puromycin incorporation, **Figure 6D**).

**Figure 6.**
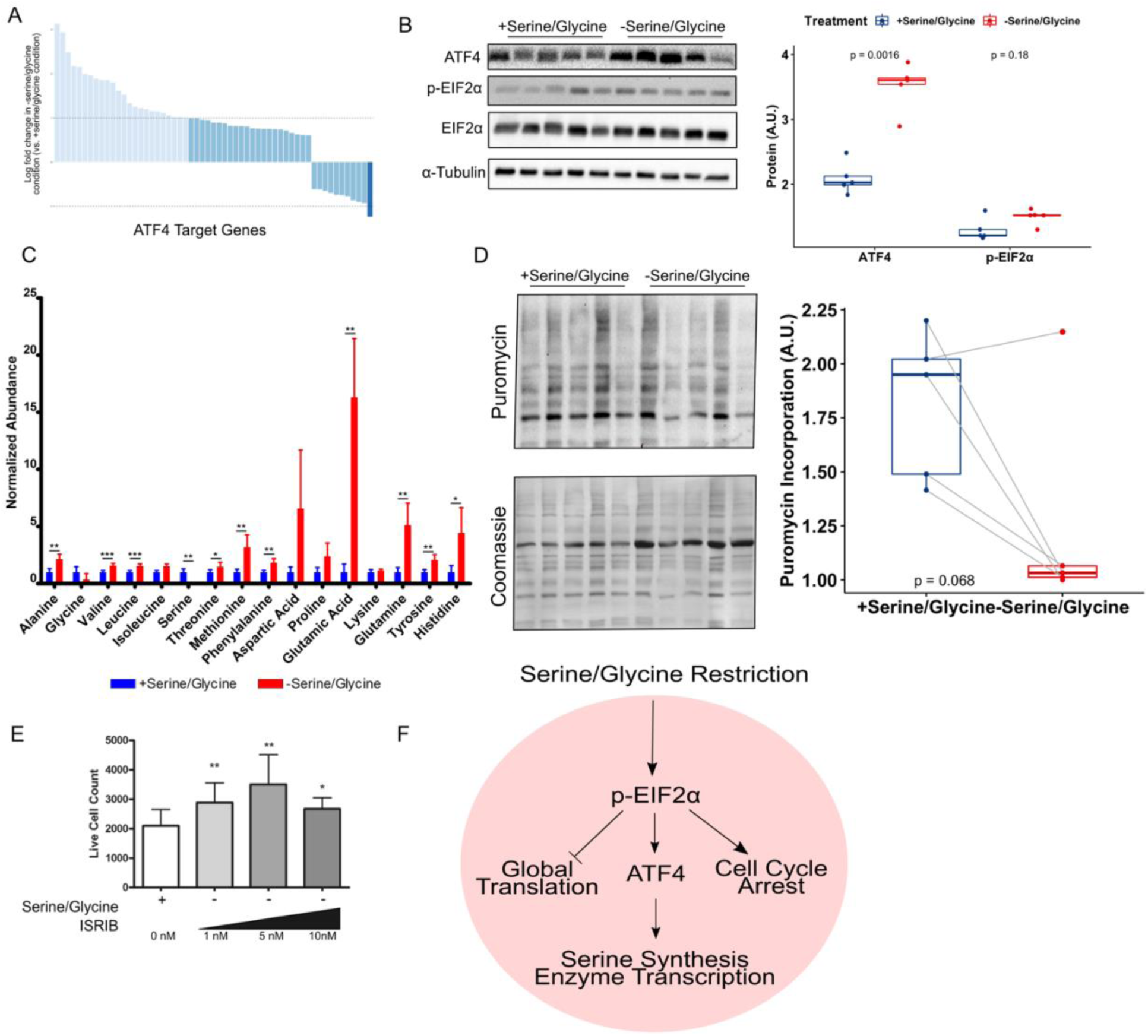
Serine/glycine restriction activates ATF4 in a p-EIFα dependent manner causing *h*MPC proliferation arrest. A) Expression of known ATF4 targets based on RNA-sequencing of *h*MPCs cultured in serine/glycine restricted media or serine/glycine replete media for 5 days. Average log fold change of transcripts in serine/glycine restricted samples vs. serine/glycine replete samples. Dark blue indicates fold change < −1 and light blue indicates fold change > 1. B) Left, protein levels of ATF4, p-eIF2α, and total eIF2α determined by immunoblotting *h*MPCs cultured in serine/glycine replete or restricted media. All proteins were normalized to α-TUBULIN expression. Right, quantification of protein expression. C) Intracellular amino acid abundance, normalized to total ion count, in *h*MPCs cultured in serine/glycine replete or restricted media for 5 days. D) Left, immunoblot analysis for puromycin incorporation of *h*MPCs cultured in serine/glycine replete or restricted media for 5 days normalized to Coomassie staining. Right, quantification of puromycin incorporation. E) Protein levels of p-eIF2α and total eIF2α determined by immunoblotting in *h*MPCs cultured in serine/glycine restricted media with and without 5 nM ISRIB. All proteins were normalized to α-TUBULIN expression. Right, quantification of protein expression. F) Live cell count after 5 days of serine/glycine restriction with or without varying doses of ISRIB. G) Generalized model of how serine/glycine restriction affects EIF2ɑ and ATF4 to reduce global protein synthesis and cause cell cycle arrest. All experiments were repeated with *h*MPCs derived from the same 5 donors. P-values are indicated on the appropriate graphs.

To directly test whether the p-eIF2α response was responsible for arrest of *h*MPC proliferation, *h*MPCs were cultured with the small molecule, p-eIF2α inhibitor, ISRIB (Sidrauski et al., 2015) (**Figure 6E**). Treatment with ISRIB increased *h*MPC cell number in the absence of serine/glycine (**Figure 6F**). Interestingly, ISRIB treatment, in the presence of serine/glycine, negatively affected cell number (**Figure S2**) suggesting that the benefit of reducing p-eIF2α signaling on *h*MPC proliferation was only effective when aberrant signaling was induced by serine/glycine restriction. Therefore, we propose that serine/glycine restriction reduces the intracellular availability of serine, glycine, and glutathione, which activates the integrated stress response and leads to limited global translation and further to cell cycle arrest (**Figure 6G**). Overall, this process is likely exacerbated in older adults who lose serine availability in the microenvironment.

## Discussion

Here, we report that aging reduces levels of serine in the *h*MPC microenvironment and that *h*MPCs are limited in their capacity for serine/glycine biosynthesis. The negative correlation between skeletal muscle levels of serine and age are in line with previous analyses comparing serum of younger and older individuals (Dunn et al., 2014; Menni et al., 2013). While glycine has not consistently been noted as reduced in the circulation of older adults, it has been shown to be reduced in the red blood cells of older adults and these reduced levels can be corrected by dietary supplementation (Sekhar et al., 2011). In support of a dysregulation of serine synthesis with age, we identified changes in expression of the serine synthesis enzymes with age in whole skeletal muscle tissue and in isolated *h*MPCs. Others have previously identified that the expression of metabolic enzymes and nutrient transporters can decline with age, however, the underlying reason for this remains elusive (Kawase et al., 2015). While our data are supportive of a decrease of serine levels in human skeletal muscle with age, potentially due to reduced biosynthesis, a direct measurement is necessary to definitively answer this question. Therefore, it is likely that reduced availability of serine/glycine to *h*MPCs is due to a combination of reduced *de novo* synthesis and dietary changes or metabolism with age.

The importance of decreased serine/glycine availability to *h*MPCs with age is highlighted by our identification of an absolute requirement for the non-essential amino acids, serine and glycine, for *h*MPC proliferation. A previous report, in rat MPCs, demonstrated that serine and glycine are required for proliferation likely due to a limited capacity for *de novo* serine synthesis (Dufresne et al., 1976). A more recent report found that C2C12 cells, an immortalized murine cell line mimicking MPCs, could proliferate in the presence of serine and absence of glycine but that proliferation was enhanced with the addition of glycine (Sun et al., 2016). It is possible that glycine in the C2C12 cell model is synthesized from serine to maintain necessary glycine for cellular processes, however, isotope tracing was not conducted to test this theory. The work presented here builds upon these previous findings by extending them into a primary, human model and elucidating the mechanism for the *h*MPC serine/glycine requirement.

The skeletal muscle of older individuals has a diminished ability to regenerate after injury due to a decline in the function of MPCs. It has been demonstrated that repeated muscle injury in aged mice depletes the number of self-renewing MPCs and consequently skeletal muscle regeneration (Sousa-Victor et al., 2014). *h*MPCs isolated from older (vs. younger) donors exhibit diminished antioxidant capacity (Fulle et al., 2005) and *h*MPCs that are undergoing replicative senescence in culture, to model aging, exhibit increased ROS levels (Minet and Gaster, 2012). Together, this suggests that *h*MPCs from older individuals may have diminished skeletal muscle regeneration because of an inability to buffer the ROS produced as a natural consequence of proliferation (L’Honoré et al., 2018). Serine/glycine are required for glutathione synthesis to maintain physiological levels of ROS (L’Honoré et al., 2018) during proliferation as demonstrated by our experiments in which supplementation of cell-permeable glutathione, increased cell number in a modest but repeatable manner. Therefore, a decline in serine availability in the *h*MPC microenvironment, coupled with a decline in the *h*MPC-intrinsic capacity for *de novo* serine synthesis, may prevent adequate glutathione synthesis to buffer proliferation associated increases in intracellular ROS and lead to a reduction in the MPC pool in older individuals. Evidence exists that dietary intervention may be able to help maintain glutathione levels with advancing age. For example, a persistent metabolic phenotype associated with aging skeletal muscle is increased ROS due to impaired glutathione synthesis and this phenotype can be corrected with dietary supplementation of the glutathione precursors glycine and cysteine (Sekhar et al., 2011). Future studies should examine the effect of dietary supplementation on glutathione production by aged MPCs.

There is a growing body of literature demonstrating that dietary glycine supplementation protects muscle mass and function during a number of disease states including cancer cachexia (Ham et al., 2014), muscle dystrophy (Ham et al., 2019), sepsis (Ham et al., 2016), and reduced calorie intake (Caldow et al., 2016). To date the beneficial effects of glycine supplementation on skeletal muscle in the disease state has been attributed to glycine’s systemic anti-inflammatory effects, its contribution to balancing ROS signaling in whole skeletal muscle tissue, and its ability to restore skeletal muscle’s anabolic response to leucine (Koopman et al., 2017). However, muscle dystrophy, specifically, is characterized by chronic MPC proliferation (Carnwath and Shotton, 1987) and genetic perturbations in MPCs have been shown to improve mouse models of muscle dystrophy by supporting maintenance of the MPC population (Gallot et al., 2018). Therefore, while much research has focused on understanding the role of dietary glycine supplementation on mature skeletal muscle as a mechanism for attenuating disease related skeletal muscle loss our data suggest that such dietary supplementation may also contribute to skeletal muscle maintenance by supporting MPC proliferation.

We demonstrate that inadequate intracellular serine/glycine and/or subsequent alterations in intracellular metabolism can be sensed in a p-eIF2α dependent manner resulting in halted protein translation and cell cycle arrest. eIF2α is a stress-sensing kinase that can integrate inputs from a number of signals to elicit the integrated stress response that directs ATF4 to upregulate genes associated with metabolism and nutrient uptake while limiting the expression of genes that are non-essential for immediate survival (Harding et al., 2003). The serine/glycine biosynthesis enzymes have been identified as ATF4 targets (DeNicola et al., 2015), highlighting the importance of this pathway in mediating cell survival under stress conditions. It is unknown whether the stress signal, which elicited the ATF4 response in *h*MPCs, was the low levels of serine/glycine themselves, the buildup of ROS caused by reduced glutathione levels, or through another mechanism. eIF2α is also a primary regulator of protein translation (Holcik and Sonenberg, 2005) and we demonstrated, in *h*MPCs, that serine/glycine restriction results in a reduction of protein translation and an accumulation of free amino acids. Inhibition of protein translation via a number of different approaches arrests cells in the G1 phase of the cell cycle (Polymenis and Aramayo, 2015). Similarly, phosphorylation of eIF2α has been previously shown to be associated with fibroblast arrest in the G1 phase of the cell cycle; however, in opposition to our results, arrest in G1 by fibroblasts was associated with a decrease in cyclin D1 translation (Hamanaka et al., 2005). In quiescent MPCs eIF2α is maintained in a phosphorylated state and ATF4 levels are abundant (Zismanov et al., 2016). When MPCs transition from activation to proliferation p-eIF2α as well as ATF4 levels decline (Zismanov et al., 2016). Furthermore, a point mutation in MPCs that prevents the phosphorylation of eIF2α forces MPCs to break quiescence, a state of cell cycle arrest, and initiate proliferation (Zismanov et al., 2016). These results are similar to those found in this report when the cell cycle arrest initiated by serine/glycine restriction was overcome by treatment with ISRIB, an inhibitor of p-eIF2α.

For the first time, we identified that *h*MPCs possess the capacity for *de novo* serine/glycine biosynthesis. Intriguingly, we note that serine/glycine biosynthesis only occurs when *h*MPCs are challenged with exogenous serine/glycine restriction. Furthermore, when serine/glycine synthesis does occur it is ineffective at restoring intracellular levels of serine/glycine to a degree necessary to support proliferation. The limited ability of *h*MPCs to produce adequate serine/glycine is at least partially attributable to a lack of NAD_+_ to facilitate the conversion of 3-phosphoglycerate to 3-phosphohydroxypyruvate and eventually serine rather than a lack of glucose to form 3-phosphoglycerate. This assertion is supported by the fact that addition of pyruvate which is converted to lactate via lactate dehydrogenase, a reaction which also oxidizes NADH to NAD_+_, increased the ratio of NAD_+_/NADH and rescued *h*MPC proliferation. Additionally, serine/glycine restriction did not increase glucose uptake nor did increasing levels of available glucose influence *h*MPC proliferation, suggesting that the limitation of serine synthesis was not driven by glycolytic substrate availability. We did note that serine/glycine restriction did not increase cell death and therefore it may be that serine/glycine biosynthesis is upregulated in serine/glycine deplete conditions to support cell survival.

In conclusion, we have outlined a novel requirement for exogenous serine/glycine to support *h*MPC proliferation. Furthermore, we have identified that the reduced availability of extracellular and intracellular serine/glycine in aging may contribute to the decline in *h*MPC-based skeletal muscle regeneration that occurs with aging.

## Methods

### Participants

Younger (21-40 years) and older adults (65-80 years) were recruited from the Tompkins County, New York area. Participants were excluded if they had a history of negative or allergic reactions to local anesthetic, used immunosuppressive medications, were prescribed to anti-coagulation therapy, were pregnant, had a musculoskeletal disorder, suffered from alcoholism (>11 drinks per week for women and >14 drinks per week for men) or other drug addictions, or were acutely ill at the time of participation (Gheller et al., 2019a; Riddle et al., 2018a). The Cornell University, Institutional Review Board approved the protocol and all subjects gave written informed consent in accordance with the Declaration of Helsinki.

### Human skeletal muscle biopsies

Skeletal muscle tissue was obtained from the vastus lateralis muscle of humans using the percutaneous biopsy technique. Visible connective or adipose tissues were removed at the time of biopsy. For tissue homogenate experiments the biopsy sample was measured for wet weight and then snap-frozen in liquid nitrogen and stored at −80°C. For cell culture experiments a 60-100 mg portion of the tissue was stored in Hibernate®-A medium (Invitrogen) at 4°C until tissue disassociation was performed (within 48 hours).

### Amino acid analysis of human skeletal muscle tissue

Frozen tissue samples (20-30 mg) were homogenized for 2 min using ceramic beads (Precellys 2 mL Hard Tissue Homogenizing Ceramic Beads Kit, Bertin Instruments, US) in 500 μL −20°C methanol, 400 μL ice-cold saline, and 100 μL ice-cold H2O containing amino acid isotope labelled internal standards (Cambridge Isotope Laboratories, #MSK-A2-1.2). An aliquot of tissue homogenate (50 μL) was dried under air and resuspended in RIPA buffer for protein quantification using bicinchoninic acid assay (BCA, BCA Protein Assay, Lambda, Biotech Inc., US). 1 mL of chloroform was added to the remaining homogenate and the samples were vortexed for 5 min followed by centrifugation at 4°C for 5 min at 15 000 g. The organic phase was collected and the remaining polar phase was re-extracted with 1 mL of chloroform. An aliquot of the polar phase was collected, vacuum-dried at 4°C, and subsequently derivatized with 2% (w/v) methoxyamine hydrochloride (Thermo Scientific) in pyridine for 60 min following by 30 min sialyation N-tertbutyldimethylsilyl-N-methyltrifluoroacetamide (MTBSTFA) with 1% tert-butyldimethylchlorosilane (tBDMS) (Regis Technologies) at 37°C. Polar derivatives were analyzed by gas chromatography (GC)-mass spectrometry (MS) using a DB-35MS column (30m x 0.25 mm i.d. x 0.25 μm, Agilent J&W Scientific) installed in an Agilent 7890A GC interfaced with an Agilent 5975C MS as previously described (Wallace et al., 2018).

### Amino acid analysis of human plasma

3 μL of human plasma were spiked with 3 μL amino acid isotope labelled internal standards (Cambridge Isotope Laboratories, #MSK-A2-1.2) and extracted with 250 μL –20 °C methanol for 10 min and centrifuged at 4°C for 10 min at 15 000 g. 200 μL of supernatant was collected, vacuum-dried at 4°C, and derivatized with MTBSTFA and tBDMS as described for muscle polar analysis.

### Primary hMPC culture

Primary *h*MPC cultures were obtained as previously described (Riddle, Bender and A. E. Thalacker-Mercer, 2018; Riddle, Bender and A. Thalacker-Mercer, 2018; Gheller, J. Blum, *et al*., 2019). Briefly, skeletal muscle tissue, stored in Hibernate®-A medium (Gibco), was minced and washed via gravity with Dulbecco’s PBS (Gibco) and then digested using mechanical and enzymatic digestion in low glucose Dulbecco’s Modified Eagle Medium (Gibco). This solution was passed through a 70 μm cell strainer into 5 mL of a growth media comprised of Ham’s F12 (Gibco), 20% FBS, 1% penicillin/streptomycin (Corning), and 5 ng/mL recombinant human basic fibroblast growth factor (bFGF, Promega) then centrifuged. The pelleted cells were resuspended in growth media containing 10% DMSO and cryopreserved at −80°C until isolation via flow cytometry. Primary *h*MPCs were sorted using fluorescence activated cell sorting with fluorescently-conjugated antibodies to cell surface antigens specific to *h*MPCs [CD56 (NCAM, BD Pharmingen) and CD29 (β1-integrin, BioLegend)] and a viability stain (7-Aminoactinomycin D, eBioscience) (Xu et al., 2015). Passage six *h*MPCs were used for all experiments and were cultured in a 5% CO_2_ atmosphere at 37°C on collagen coated plates (Type I, Rat Tail, Corning). For cell culture experiments, donor cells from females were used exclusively due to availability of adequate sample.

*h*MPCs were initially seeded in the growth medium described above before being switched to treatment medium 24 hours later. For all experiments a specially formulated DMEM devoid of serine, glycine, methionine, choline, pyridoxine, glucose, folate, nucleotides, and nucleosides that was supplemented with dialyzed and charcoal treated FBS (10%), 200 µM methionine, 4 mg/L pyridoxine, 5 ng/mL bFGF, 25 nM (6S) 5-formylTHF, penicillin-streptomycin (1%), 10 mmol/L glucose, 4 mM Glutamax (Gibco), as well as L-serine and glycine in varying concentrations. Unless otherwise noted, all assays were performed after 5 days in the treatment medium specified as this was the time point when differences in cell number between cells grown in serine/glycine restricted and replete media were initially identified. In all experiments cell culture media was replenished daily. All experiments contained appropriate vehicle controls (DMSO or sterile H_2_O).

### RNA isolation and quantification

For quantitative RT-PCR analysis and RNA-sequencing (RNA-seq), RNA was isolated from *h*MPCs and using Omega E.Z.N.A.® Total RNA Kit I (Omega) according to the manufacturer’s instructions. RNA was isolated from skeletal muscle biopsy tissue using Trizol Reagent (Ambion) as per manufacturer’s instructions. RNA quantity was determined spectrophotometrically.

### Quantitative RT-PCR

Gene expression was measured using quantitative RT-PCR. cDNA was synthesized via reverse transcription of extracted RNA using the Applied Biosystems High-Capacity cDNA Reverse Transcription Kit. The Taqman Gene Expression System (Applied Biosystems) was used to measure mRNA expression levels of phosphoglycerate dehydrogenase (*PHGDH*, HS00358823), phosphoserine aminotransferase 1 (*PSAT1*, Hs00268565), and phosphoserine phosphatase (*PSPH*, Hs01921296). All samples were normalized to *18S* (Hs99999901) expression.

### RNA library preparation and sequencing

Prior to RNA-seq, RNA quality was determined using an AATI Fragment Analyzer; all samples had an RNA quality number >8.5. The NEBNext Ultra II RNA Library Prep Kit (New England Biolabs) was used to generate TruSeq-barcoded RNA-Seq libraries. Libraries were quantified with the Qubit 2.0 (dsDNA HS kit; Thermo Fisher) and size distribution was measured with a Fragment Analyzer (Advanced Analytical) before pooling. A NextSeq500 (Illumina) was used for sequencing, and a minimum of 20 M single-end 75 bp reads per library were obtained. Cutadapt v1.8 was used to trim low quality reads and adaptor sequences (parameters: -m 50 -q 20 -a AGATCGGAAGAGCACACGTCTGAACTCCAG–match-read-wild cards) (Martin, 2011). Tophat v2.1 was used to map reads to the reference genome (parameters:-library-type=fr-firststrandon-no-novel-juncs-G<ref_genes.gtf>) (Kim et al., 2013). Differential gene expression analysis was performed using *edgeR* (McCarthy et al., 2012). Only genes with at least 2 counts per million in at least three of the samples were retained for analysis.

### Immunoblot analysis

For immunoblot analysis, protein was isolated from *h*MPCs with RIPA buffer containing phosphatase (PhosSTOP, Roche) and protease (cOmplete, Roche) inhibitors. The protein concentration was quantified by BCA. 8-15 μg of protein was loaded on 10% SDS gels and transferred to PVDF membranes. Membranes were incubated in primary antibodies PHGDH (Proteintech), PSAT1 (Proteintech), PSPH (Proteintech), SHMT1 (Stover laboratory) (Woeller et al., 2007), SHMT2 (Stover laboratory) (Anderson and Stover, 2009), Cyclin D1 (Cell Signaling), ATF4 (Cell Signaling), phosphorylated (Ser51) p-EIF2α (Cell Signaling), and total EIF2α (Cell Signaling) diluted (1:1000) in a chemiluminescent blocking buffer (bl∅kTM – CH, Millipore) overnight at 4oC. After the overnight incubation, membranes were washed with 0.1% Tween in tris-buffered saline before incubation with appropriate secondary antibody (rabbit, Proteintech; goat, Thermo Scientific; mouse, Proteintech; goat, Piece) at a 1:100000 dilution in a chemiluminescent blocking buffer, at room temperature, for 60 min. Membranes were visualized after a brief incubation in SuperSignalTM West Femto (Thermo Scientific) on the Bio-Rad ChemiDoc MP. Protein expression was normalized to α-TUBULIN expression (Cell Signaling) using the ImageLab 4.1 software (Bio-Rad).

### Live and dead cell counting

Live cell number was determined by co-staining cells with Hoechst 33342 (to identify number of nuclei, Life Technologies) and propidium iodide (to identify dead cells, ThermoFisher Scientific). The number of live cells was determined by subtracting the number of propidium iodide positive cells from the Hoechst 33342 positive cells as identified using the Celigo imaging cytometer (Nexcelcom).

### Cell cycle analysis

For cell cycle analysis, *h*MPCs were pelleted, washed with ice-cold PBS, re-pelleted and resuspended in PBS then fixed in 3:1 volume:volume 100% ice-cold ethanol before being stored at 4 °C overnight. The following day, cells were pelleted, washed with ice-cold PBS, and resuspended in 400 µL PBS containing 2 mM EDTA. Cell suspensions were incubated with 100 µL 10 mg/mL RNAse A (VWR) for 30 min at 37oC, to degrade RNA, followed by DNA staining with 50 µL of 1 mg/mL propidium iodide for 30 min, in the dark, at room temperature. Propidium iodide intensity was measured using a flow cytometer (BD Aria Fusion). The percentage of the total population of cells in G1, S-phase, G2 cells were determined using FlowJo’s (Becton, Dickinson, and Company) univariate platform.

### 5-bromo-2’-deoxyuridine (BrdU) incorporation

To determine the proportion of cells actively synthesizing DNA, *h*MPCs were pulsed with BrdU for 24 hours. *h*MPCs were washed with prewarmed PBS before being fixed in ice-cold methanol for 5 min. Cells were washed with PBS before 30 min of acid hydrolysis and cell-permeabilization in 2 N HCl prepared in 0.1% PBS-Tween20. Cells were washed with PBS before being blocked in 1% BSA/10% normal goat serum/0.3 M glycine/0.1% PBS-Tween20 followed by washing in PBS. Fixed and blocked *h*MPCs were incubated overnight at 4°C with an anti-BrdU antibody (1:400 dilution, Biolegend) followed by PBS washes and a 60 min incubation with an Alexa-Fluor 488 conjugated anti-mouse secondary antibody (Invitrogen). Finally, *h*MPCs were washed with PBS and incubated with Hoechst 33342 before visualization and analysis using the Celigo imaging cytometer (Nexcelcom).

### Glucose uptake

Glucose uptake was measured based on the detection of 2-deoxyglucose-6-phosphate uptake by a commercially available luminescence-based kit (Glucose Uptake-GloTM Assay, Promega) on a SpectraMax M3 (Molecular Devices). Values were normalized to total cell count obtained from a parallel plate.

### Stable-Isotope Labeling, Metabolite Extraction, and GC-MS Analysis

For isotopic labeling experiments, cells were cultured in 10 mM [U-_13_C_6_]glucose (Cambridge Isotope Laboratories, Inc.) containing either 1000 µM serine/glycine or no serine/glycine for 48 h prior to extraction. A medium exchange was performed after 24 hours. On the day of extraction, polar metabolites were extracted as previously described (Cordes and Metallo, 2019).

The upper aqueous phase was derivatized using a Gerstel MPS with 15 µL of 2% (w/v) methoxyamine hydrochloride (Thermo Scientific) in pyridine (incubated for 60 min at 45°C) and followed by 15 µL MTBSTFA with 1% tert-butyldimethylchlorosilane (Regis Technologies) (incubated for 30 min at 45°C). Polar derivatives were analyzed by GC-MS using a DB-35MS column (30 m x 0.25 mm i.d. x 0.25 µm) installed in an Agilent 7890B GC interfaced with an Agilent 5977B MS with an XTR EI source using the following temperature program: 100°C initial, increase by 3.5°C/min to 255°C, increase by 15°C/min to 320°C and hold for 3 min.

### Glutathione measurements

Total and oxidized (GSSG) glutathione were measured using a commercially available luminescence-based kit (GSH-GSSG GloTM Assay, Promega) on a SpectraMax M3 (Molecular Devices). The reduced (GSH) to oxidized GSSG ratio was determined by multiplying the GSSG reading by 2, to account for each mole of oxidized GSSG producing two moles of total glutathione, subtracting that number from the total glutathione levels and finally, dividing this value by the total GSSG reading. Values were normalized to total cell count obtained from a parallel plate.

### Reactive oxygen species detection

To determine the intracellular level of ROS, *h*MPCs were pelleted and resuspended in 1 mL pre-warmed PBS with CellROX Green Reagent (Invitrogen), a cell-permeable dye which fluoresces green and binds to DNA upon oxidation, at a final concentration of 5 µM. After a 30 min incubation at 37oC, *h*MPCs were washed with PBS before being fixed in 2% paraformaldehyde for 10 min at room temperature. *h*MPCs were again washed with PBS before finally being resuspended in 300 µL 0.5 mM EDTA in PBS and analyzed via flow cytometry (BD Aria Fusion).

### NAD/NADH measurements

The reduced NAD (NADH) to oxidized NAD (NAD_+_) ratio was determined in *h*MPCs via a commercially available luminescence-based kit (NAD/NADH GloTM Assay, Promega) following the manufacturer’s instructions. Luminescence values were obtained with a SpectraMax M3 (Molecular Devices) and normalized to total cell count obtained from a parallel plate.

### Protein synthesis

The SUnSET Method (Schmidt et al., 2009) was used to quantify the rate of puromycin incorporation to approximate protein synthesis (Henrich, 2016). *h*MPCs were treated with 0.5 µg/mL puromycin (Thermo Fisher) for 30 min and then immediately harvested in RIPA buffer and protein was isolated as described above. Immunoblotting was performed, as described, using a primary antibody specific for puromycin (Millipore) diluted 1:1000 in a chemiluminescent blocking buffer overnight, prior to incubation in secondary antibody (mouse, Proteintech) diluted 1:100000. *h*MPCs were incubated in SuperSignalTM West Femto (Thermo Scientific) and imaged on the Bio-Rad ChemiDoc MP. Puromycin expression was normalized to the total protein level in each respective lane as determined by Coomassie staining and imaging on the Bio-Rad ChemiDoc MP using the ImageLab 4.1 software (Bio-Rad).

### Statistics

Statistical analyses were performed in R Studio (Version 1.0.136). For metabolite analysis of whole skeletal muscle tissue, the normalcy of the distribution of each amino acid was assessed by the Shapiro-Wilk test. If data were determined to be normally distributed, they were compared via an unpaired t-test otherwise they were compared by a Mann-Whitney U-test. The correlation between skeletal muscle serine levels and age were determine using a Pearson correlation coefficient. When comparing gene and protein expression between age groups, an unpaired t-test was performed. For cell counting experiments, two-way analysis of variance (ANOVA) was performed with time and treatment being the main factors. A Tukey *post hoc* test was performed if the interaction term was significant (P<0.05). For all *h*MPC assays, either a paired t-test or a repeated measures ANOVA was employed.

## Acknowledgments

This work was financially supported by a President’s Council of Cornell Women Award (to A.T.M), Cornell University Division of Nutritional Sciences funds (to A.T.M), and a Canadian Institutes for Health Research Doctoral Foreign Study Award (to B.J.G), and NIH grant R01CA234245 (to C.M.M.).

## Author Contributions

Conceptualization, B.J.G, P.J.S, and A.E.T..; Methodology, B.J.G, M.S.F., P.J.S., and A.E.T..; Investigation, B.J.G, J.E.B, M.E.G., E.W.L., and M.K.H.; Resources, E.B., P.J.S, M.S.F., B.D.C., C.M., and A.E.T.; Writing – Original Draft, B.J.G. and A.E.T.; Writing – Review and Editing, B.J.G., J.E.B, M.E.G., E.W.L., M.K.H., P.J.S., M.S.F., B.D.C., C.M.M., and A.E.T.; Supervision, A.E.T.

---

**Figure S1.**
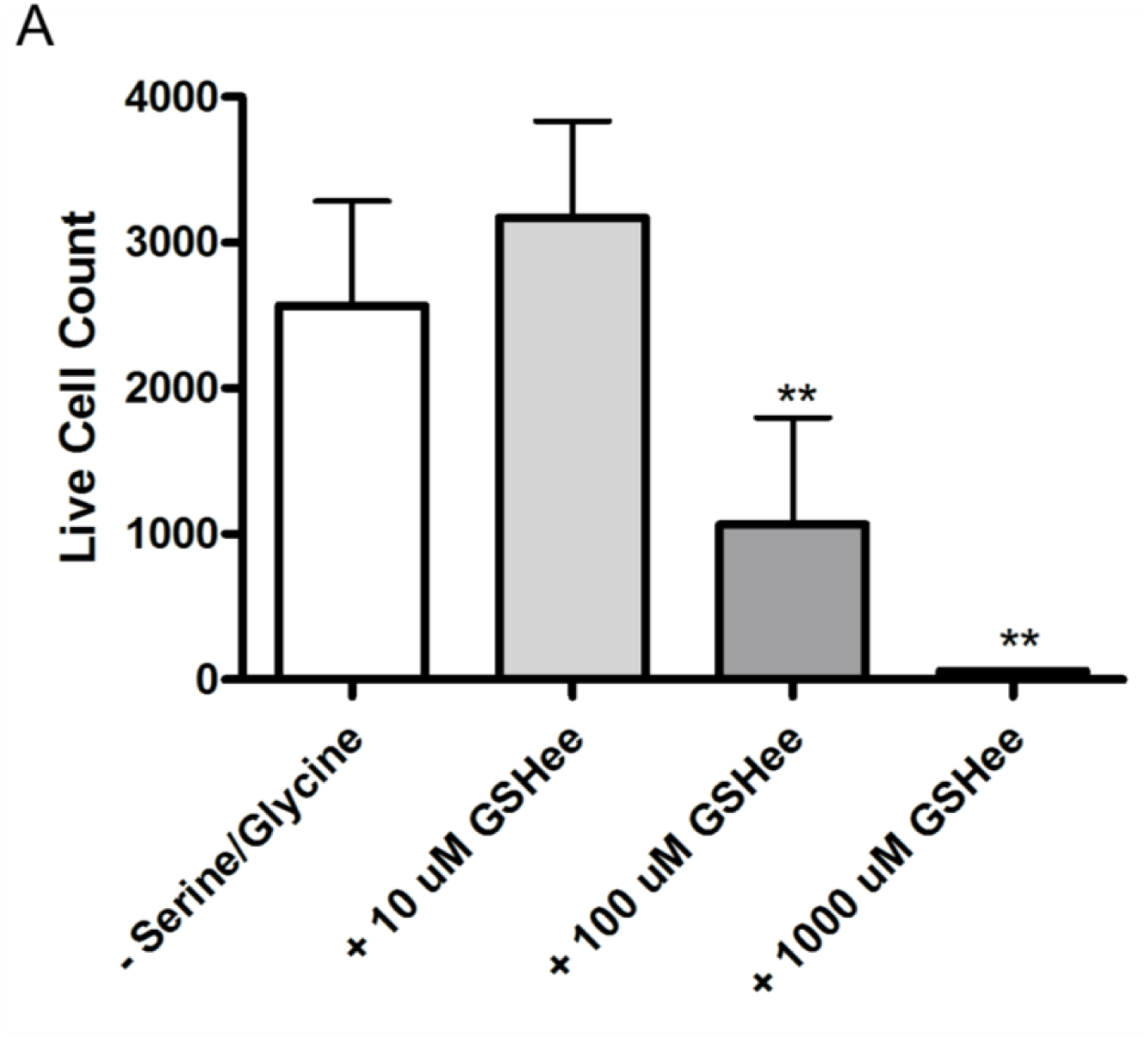
Glutathione ethyl ester is toxic to *h*MPCs at high doses A) Live cell count after 5 days of GSHee (glutathione ethyl ester) supplementation in serine/glycine restricted media. All experiments were repeated with *h*MPCs derived from the same 5 donors. **P<0.01 relative to serine/glycine restricted control. Data are expressed as mean ± SD.

**Figure S2.**
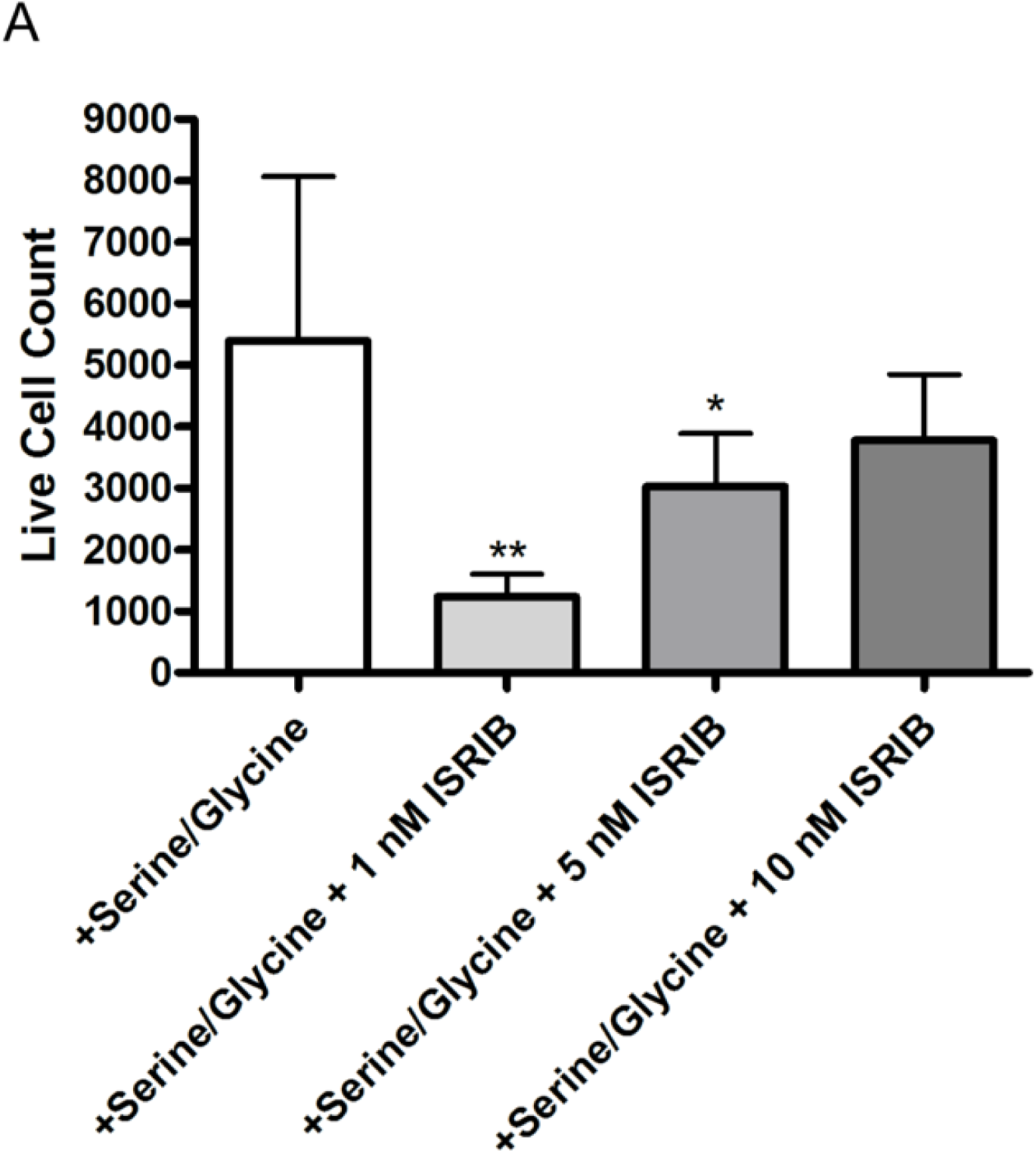
ISRIB reduces *h*MPC number in the presence of serine/glycine A) Live cell count after 5 days of ISRIB supplementation in serine/glycine replete media. All experiments were repeated with *h*MPCs derived from the same 5 donors. **P<0.01, **P<0.01 relative to serine/glycine replete control. Data are expressed as mean ± SD.

## Supplementary Tables

**Supplementary Table 1.** List of Differentially Expressed Genes in *h*MPCs Cultured in Serine/Glycine Replete and Serine/Glycine Restricted Media

